# Ovarian carcinosarcoma genomics and pre-clinical models highlight the N-MYC pathway as a key driver and susceptibility to EMT-targeting therapy

**DOI:** 10.1101/2020.11.24.396796

**Authors:** Gwo Yaw Ho, Elizabeth L. Kyran, Justin Bedo, Matthew J. Wakefield, Darren P. Ennis, Hasan B. Mirza, Elizabeth Lieschke, Cassandra J. Vandenberg, Olga Kondrashova, Rosie Upstill-Goddard, Ulla-Maja Bailey, Suzanne. Dowson, Patricia Roxburgh, Rosalind M. Glasspool, Gareth Bryson, Andrew V. Biankin, Susanna L. Cooke, on behalf of the Scottish Genomes Partnership, Gayanie Ratnayake, Orla McNally, Nadia Traficante, Australian Ovarian Cancer Study, Anna DeFazio, John Weroha, David D. Bowtell, Iain A. McNeish, Anthony T. Papenfuss, Clare L. Scott, Holly E. Barker

## Abstract

Ovarian carcinosarcoma (OCS) is an aggressive and rare tumour type with limited treatment options. OCS is hypothesised to develop via the combination theory from a single progenitor, resulting in carcinomatous and sarcomatous components, or alternatively via the conversion theory, with the sarcomatous component developing from the carcinomatous component through epithelial-to-mesenchymal transition (EMT). We show OCS from 18 women to be monoclonal through analysis of DNA variants from isolated carcinoma and sarcoma components. RNA sequencing indicated the carcinoma components were more mesenchymal when compared with pure ovarian carcinomas, supporting the conversion theory. We used pre-clinical OCS models to test the efficacy of microtubule-targeting drugs, including eribulin, which has been shown to reverse EMT characteristics. We demonstrated that microtubule inhibitors, vinorelbine and eribulin, were more effective than standard-of-care platinum-based chemotherapy. Eribulin reduced mesenchymal characteristics, N-MYC expression and cholesterol biosynthesis. Finally, eribulin induced a strong immune response, supporting immunotherapy combinations in the clinic.

## Introduction

Ovarian carcinosarcoma (OCS), also known as malignant mixed Müllerian tumour, is a heterogeneous cancer with poor prognosis^1^, accounting for 1-4% of ovarian malignancies^2,3^. These tumours contain both epithelial (carcinoma) and mesenchymal (sarcoma) components^3^. Molecular analysis suggests that most OCS are monoclonal^4–9^, with two theories for OCS histogenesis: the combination theory, where a single stem cell differentiates early to form the two components; and the conversion theory, where the carcinoma undergoes epithelial-to-mesenchymal transition (EMT) to form the sarcomatous component^10^.

*TP53* mutations and loss of heterozygosity (LOH) of 17p, and consequent chromosomal instability, are common in OCS^7,8,11–14^. Mutations in *PIK3CA*, *PTEN*, *KRAS*, *FBXW7*, *CTNNB1*, and *RB1* are observed frequently^5,8,9,13,15–17^, whilst mutations in *ARID1A*, *ARID1B, KMT2D*, *BAZ1A*, *BRCA1*, *BRCA2*, and *RAD51C* have also been reported^8,15–19.^ One study also identified recurrent mutations in the genes encoding histones H2A and H2B (*HIST1H2AB*/*C*, *HIST1H2BB*/*G*/*J*) that play a role in EMT^9^. Only one study has analysed gene expression in the separate components, finding a strong positive correlation of EMT score with sarcoma content as well as methylation of the EMT-suppressing miRNAs miR-141/200a/200b/200c/429^8^.

EMT can be induced through aberrant expression of the high-mobility-group AT-hook protein 2 (HMGA2) and subsequent activation of the TGFβ signalling pathway^20^. HMGA2 binds preferentially to AT-rich DNA sequences in a histone-independent manner^21–24^. HMGA2 is not expressed in most adult tissues^25,26^, but high expression has been observed in many cancers and is correlated with metastasis and chemotherapy resistance^27–31^. HMGA2 expression is thought to be largely controlled by the microRNA *let-7*^32–35^. Other downstream target genes of *let-7* include *MYCN* and *LIN28B*, whilst LIN28B inhibits maturation of *let-7*^36^, reinforcing both low and high expression states and acting as a bistable switch. Up-regulation of the N-MYC/LIN28B pathway has been observed in the aggressive C5 subset of ovarian or fallopian tube high-grade serous carcinoma (HGSC) and in other aggressive cancer subtypes, and is indicative of poor prognosis^36–38^. Furthermore, high HMGA2 expression has been observed in 60% of OCS cases^39^. We hypothesised that up-regulation of the N-MYC/LIN28B pathway and subsequent expression of HMGA2 may be a key driver of OCS, and drugs that target EMT may be effective.

Eribulin is a microtubule-targeting drug that binds to the plus (β tubulin exposed) end of microtubules resulting in mitotic blockade^40,41.^ *In vitro*, *in vivo* and human studies show that eribulin can reverse EMT, leading to favourable intra-tumoral vascular remodelling, reduced cell invasion, increased cell differentiation^42–46^ and modulation of the tumour-immune microenvironment^47^. Eribulin has completed Phase III trials for metastatic breast cancer, soft-tissue sarcoma and non-small cell lung cancer (NSCLC). It has Therapeutic Goods Administration (TGA) approval for treatment of advanced breast cancer and liposarcoma^47,48^. We hypothesised that eribulin may be effective against OCS tumours due to its role in reversal of EMT characteristics.

Here we present mutation, copy number and gene expression analyses of separate components from an OCS cohort. We have used a unique genetically engineered mouse model (GEMM) and patient-derived xenograft (PDX) models of OCS to assess the efficacy of a range of microtubule-targeting drugs and to determine the mechanism of action of eribulin, a drug with significant activity in these models.

## Results

### Mutation and copy number profile of OCS was similar to HGSC

We identified eighteen women diagnosed with OCS, seventeen with high-grade serous carcinoma (HGSC) and one with grade 2 endometrioid histology in the carcinoma component. Twelve associated metastatic samples were also available. Full clinical details are shown in Supplementary Table S1 and Supplementary Figure S1. Targeted sequencing of 377 genes in macro-dissected carcinoma and sarcoma components as well as metastases was performed (Supplementary Tables S2-S7; Supplementary Figure S2).

Overall, OCS samples had genomic profiles similar to HGSC, with near-ubiquitous *TP53* mutation (17/18 cases, including 17/17 with HGSC pathology), *CCNE1* amplification (4/18 cases), *BRCA2* loss or mutation (4/18 cases), *KRAS* mutation and amplification (4/18 cases), *PIK3CA* mutation and amplification (4/18 cases), *NF1* or *CDKN2A* mutation or disruption by rearrangement (2/18 cases each), *RB1* deletion (2/18 cases), *PTEN* mutation (2/18 cases) and *MYC* or *MYCN* amplification (1/18 and 2/18 cases, respectively) (Figure 1a). Overall mutational burden was low (mean 1.2, median 0.87 mutations/MB sequenced), which did not differ between carcinoma and sarcoma (Figure 1b, Supplementary Table S8). However, as with HGSC, the genomes were structurally unstable with an average of 3.3 high-level gains and 1.4 likely homozygous deletions called per sample (Supplementary Figure S3).

**Figure 1:**
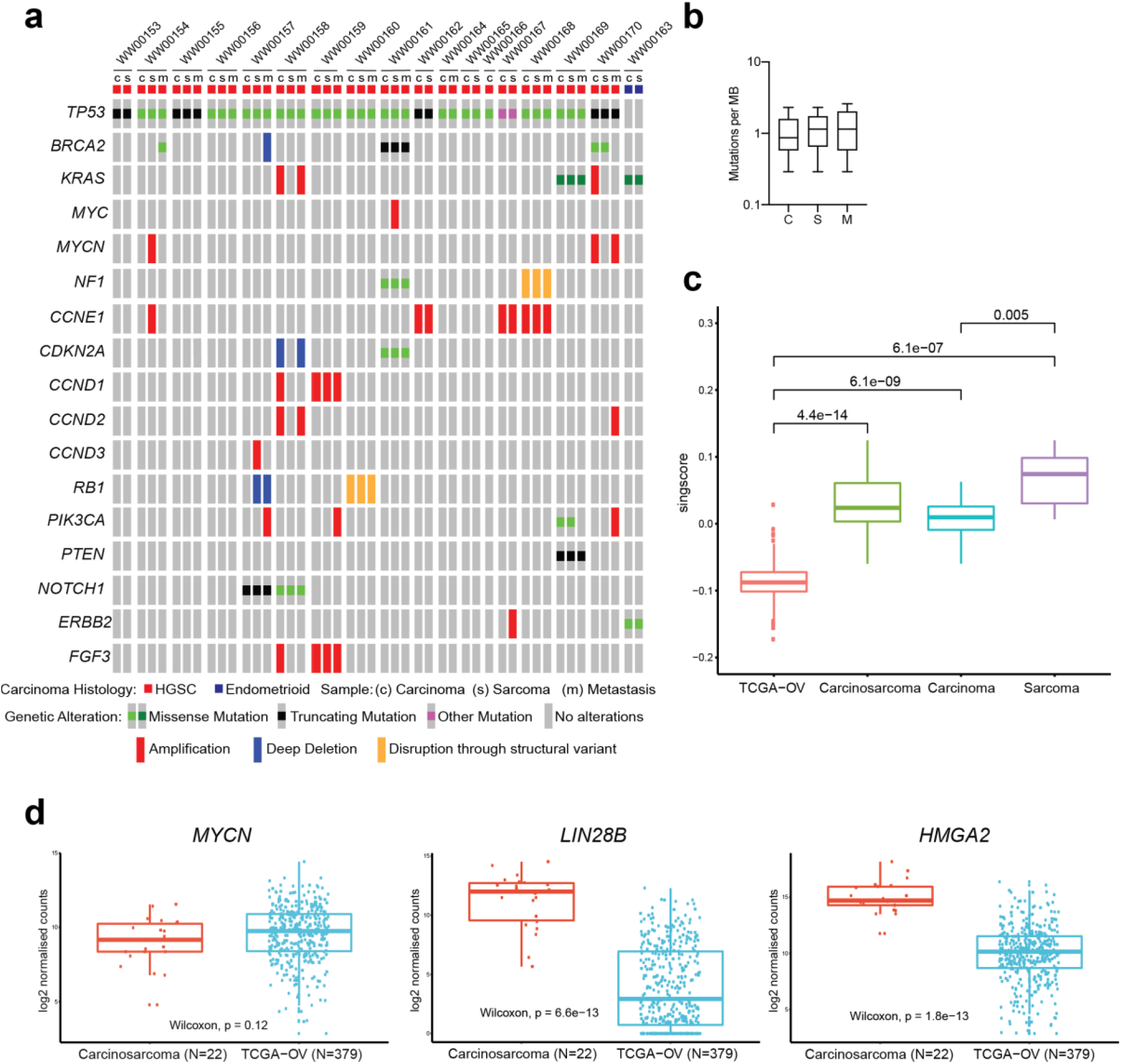
Mutational and structural variant landscape of ovarian carcinosarcoma. **(A)** Summary of frequently altered genes across the carcinoma, sarcoma and metastasis samples from 18 macrodissected ovarian carcinosarcoma samples. For missense mutations, light green represents “unknown significance” and dark green represents “putative driver”. **(B)** Mutation burden (mutations per megabase sequenced. **(C)** Comparison of EMT scores in separated carcinomatous and sarcomatous regions from ovarian carcinosarcoma samples, whole ovarian carcinosarcoma tumours, and ovarian high-grade serous carcinoma samples in TCGA. **(D)** Expression of *MYCN*, *LIN28B* and *HMGA2* in our ovarian carcinosarcoma cohort compared to ovarian high-grade serous carcinoma tumours in TCGA. TCGA-OV, ovarian high-grade serous carcinomas in TCGA; C, carcinoma; S, sarcoma; M, metastasis.

Only WW00163 lacked a *TP53* mutation. It had mutations in *KRAS* and *ERBB2* (Figure 1a) together with a subclonal mutation of *KMT2C* and lacked the genomic chaos typical of HGSC (Supplementary Figure S4), in keeping with an origin of endometrioid carcinoma.

Based on point mutation profiles, there were no consistent differences between the sarcoma and carcinoma components. In all cases, the two components shared at least one point mutation, demonstrating a shared clonal origin. Half of carcinoma-sarcoma pairs (8/16) shared all point mutations while the others gained additional mutation(s) in one or both components. On average, carcinoma-sarcoma pairs differed by only a single mutation (range 0-7). These data indicate that these tumours are monoclonal, which supports both the conversion and combination theories of carcinogenesis.

By contrast, there were more copy number changes between the carcinoma and sarcoma components, with an average of 10.6 genes having a different copy number state between the two (range 0-36) (Supplementary Figures S3 and S4; Supplementary Table S6). The most commonly different genes were *FGF3* and *MDM2* (Supplementary Table S7). However, these differences did not appear to be focal or high level, perhaps suggesting that these genes are not specific targets of alteration between carcinomas and sarcomas. Instead these chromosomal differences may arise due to ongoing chromosomal instability. Case WW00169 had neither mutation nor copy number differences between the carcinoma and sarcoma components.

Interestingly, in some cases metastases showed substantial genomic divergence from their corresponding primary, indicative of an early seeding to the metastatic sites (Figure 1a). In addition to two cases (WW00154, WW00158) where the metastasis either gained three mutations or lost four, a third case (WW00157) diverged in several likely driver copy number events including loss of *BRCA2* between the carcinoma and its corresponding metastasis (Supplementary Tables S4, S6 and S7).

### OCS had EMT-like and N-MYC pathway gene expression patterns

We next undertook RNA sequencing (RNAseq) on isolated carcinoma (n=13) and sarcoma (n=9, 7 paired with carcinoma) components (Supplementary Figure S5; Supplementary Tables S9-S12).

Using an EMT expression signature derived from uterine carcinosarcoma^49^, we found a highly significant enrichment of EMT in carcinosarcomas, compared with the TCGA cohort of ovarian HGSC (TCGA-OV; n=379)^50^. This enrichment was predominantly driven by the sarcoma component (*p*<0.0001; Figure 1c) and was confirmed using other reported EMT signatures^51–53^ (Supplementary Figure S6). Interestingly, the carcinoma components also had significantly higher EMT scores than the TCGA-OV cohort, suggesting that the OCS carcinoma component was either predisposed to undergo sarcomatous transformation or already transitioning to sarcoma (*p*<0.0001; Figure 1c). Together, these data support the conversion theory of OCS development.

To study the N-MYC/LIN28B pathway specifically, we analysed *MYCN*, *LIN28B* and *HMGA2* expression in the same dataset. *LIN28B* and *HMGA2* were significantly up-regulated compared to the TCGA-OV cohort (*p*<0.0001 for both; Figure 1d).

### p53 inhibition and up-regulation of the N-MYC/LIN28B pathway in fallopian tube secretory epithelial cells gave rise to OCS

We established an OCS GEMM by directing both p53 inhibition and N-MYC/LIN28B pathway up-regulation to the fallopian tube secretory epithelial cell (FTSEC) via the PAX8 promoter^54–57^. The resulting founder tumour (T0) and stable cell line derived from a first passage tumour (T1) (OCS GEMM cells) were used for subsequent experiments (Figure 2a; Supplementary Tables S13 and S14).

**Figure 2:**
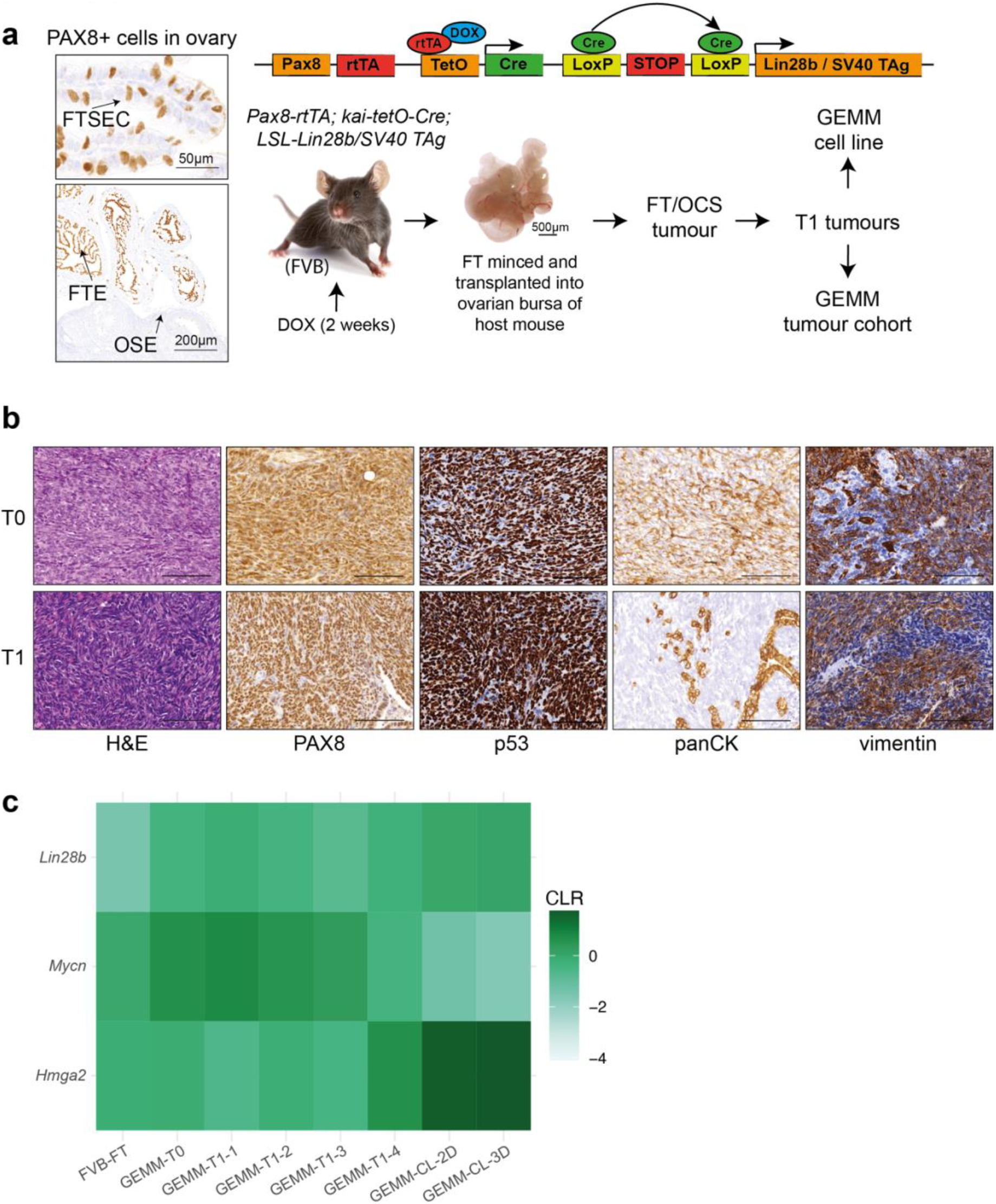
Generation and characterisation of a GEMM of OCS. **(A)** A *Pax8-rtTA*; *kai-tetO-Cre*; *LSL-Lin28b/SV40TAg* transgenic mouse was treated with doxycycline for 2 weeks to induce expression of Cre recombinase in the FTSECs. The FTs were then removed, minced and transplanted into the ovarian bursa of a cba/Nu host mouse, generating the GEMM founder tumour (T0). This tumour was transplanted into mice to establish the first and subsequent OCS cohorts (T1, T2, T3, etc). A T1 tumour was also digested and cultured *in vitro* to generate a cell line. **(B)** The GEMM T0 founder tumour and a tumour generated after the 1^st^ transplant (T1) were assessed by IHC. T0 and T1 tumours expressed PAX8, indicating FTSEC derivation. Representative images of H&E, PAX8, p53, Pan-CK and Vimentin staining are shown. Scale bars represent 100μm. **(C)** The GEMM T0 founder tumour, T1 tumours (n=4) and the GEMM cell line grown in 2D and 3D were analysed by RNA-seq. Controls included normal FT tissue harvested from FVB mice, and FT epithelial cells and stroma. A heatmap shows the expression of genes involved in the N-MYC/LIN28B pathway: *Lin28b*, *Mycn* and *Hmga2*. GEMM, genetically engineered mouse model; FTSEC, fallopian tube secretory epithelial cell; FTE, fallopian tube epithelium; OSE, ovarian surface epithelium; CK, cytokeratin; FT, fallopian tube; CLR, centred log ratio.

IHC analysis revealed high p53 expression, in keeping with SV40 TAg-mediated accumulation^58^ (Figure 2b). Tumours expressed cytokeratin (pan-CK) in approximately 5% and vimentin in approximately 95% of the regions analysed, indicating a predominantly sarcomatous phenotype (Figure 2b). RNA sequencing confirmed up-regulation of *Lin28b* and *Mycn* in the tumours and up-regulation of *Lin28b* and *Hmga2* in the cell line, relative to control fallopian tubes (Figure 2c; Supplementary Table S15), whilst quantitative RT-PCR confirmed elevated expression of *Lin28b* in both the tumour and cell line (Supplementary Figure S7).

### GEMM tumours were resistant to current standard-of-care treatments but responded to the microtubule inhibitors vinorelbine and eribulin

We assessed the *in vivo* response of GEMM tumours to standard-of-care HGSC therapies; cisplatin, pegylated liposomal doxorubicin (PLD) and paclitaxel. Overall, the tumours were refractory to all three treatments, as the time to progressive disease (PD) was the same as for vehicle treatment. PLD and cisplatin failed to demonstrate any meaningful response in the GEMM tumours (Figure 3a), although paclitaxel demonstrated modest responses with an increase in median time-to-harvest (TTH) from 15 to 36 days compared to vehicle treatment (Table 1, *p*=0.0101, respectively). By contrast, significant tumour regression was observed in all tumours treated with the microtubule inhibitor vinorelbine leading to improvement of median TTH (15 days (vehicle) vs 81 days (vinorelbine); Figure 3a, Table 1; *p*<0.0001). Eribulin also resulted in significant tumour regression in all tumours leading to improvement of median TTH (15 days (vehicle) vs 46 days (eribulin); Figure 3a, Table 1; *p*<0.0001). Expression of Ki67 in the tumours was reduced one week after mice received a single dose of eribulin (Figure 3b).

**Figure 3:**
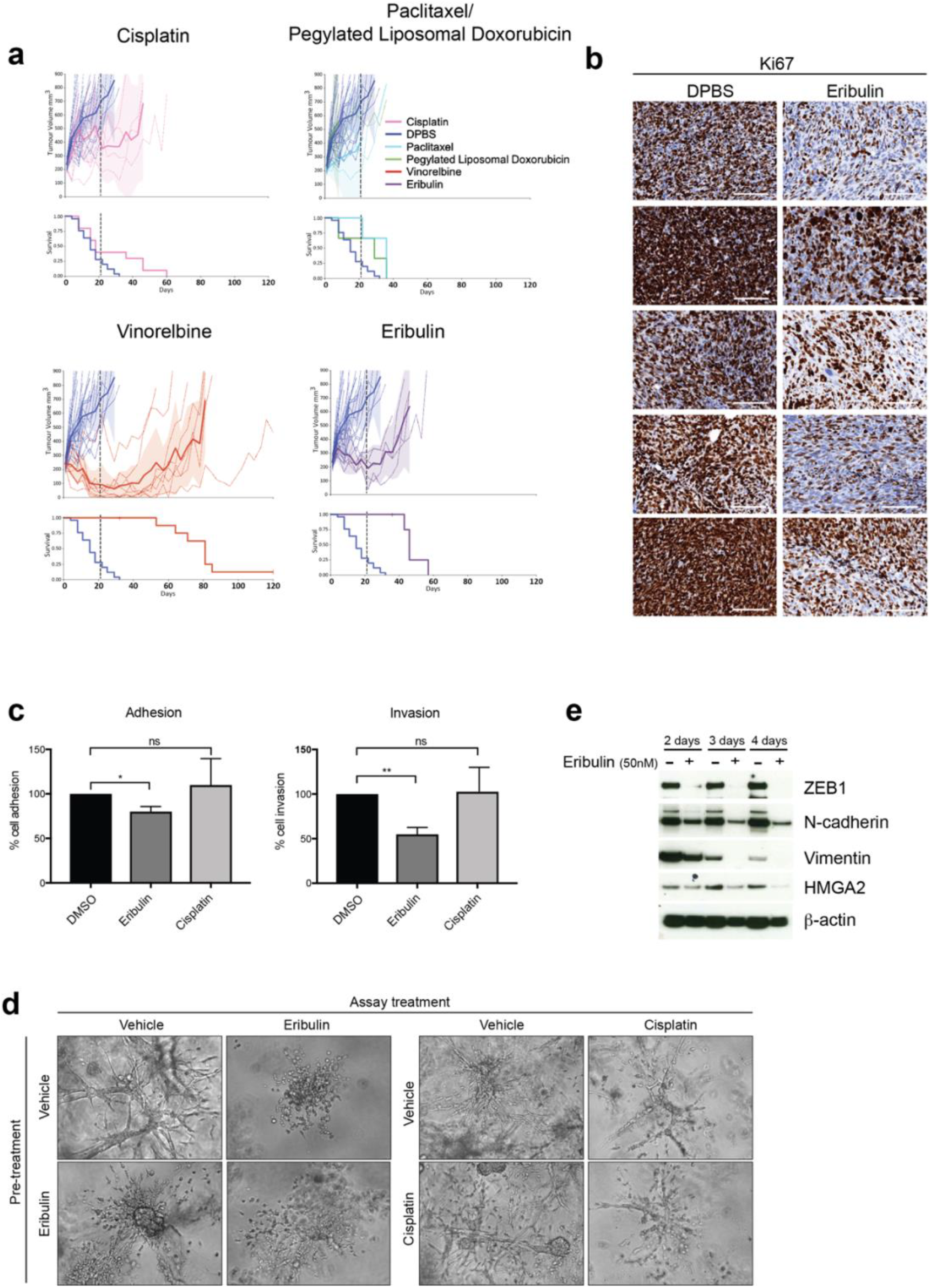
GEMM OCS tumours are refractory to current standard-of-care treatments for ovarian cancer but are responsive to the microtubule drugs vinorelbine and eribulin. **(A)** *In vivo* treatment of GEMM OCS tumours with: DPBS (n=25), cisplatin (4mg/kg; n=10), PLD (1.5mg/kg; n=3), paclitaxel (n=3), vinorelbine (15mg/kg; n = 9) and eribulin (1.5mg/kg; n=5). Shaded area = 95% confidence interval. Time to PD and harvest (TTH) are shown in Table 1. **(B)** Expression of Ki67 was assessed by IHC in a number of tumours after a single dose of eribulin (or DPBS vehicle). Representative images are shown. Scale bars represent 100μm. **(C)** GEMM cells were pre-treated with eribulin (20 nM), cisplatin (0.2 μM) or vehicle control (DMSO) for one week before being plated in adhesion assays (left panel) or migration and invasion assays (right panel). Percentage of adherent cells was calculated compared to vehicle-treated controls. Percentage of invading cells was calculated compared to number of migrating cells. **(D)** GEMM cells were pre-treated as above with eribulin, cisplatin or vehicle control (DMSO) for one week before being plated in collagen with treatment either removed or maintained. Representative images of colonies growing in collagen on day 8 are shown. Scale bars represent 200μm. **(E)** Expression of the mesenchymal markers ZEB1, N-cadherin, Vimentin and HMGA2 in cells exposed to 50nM eribulin or DMSO control for the indicated time-points was determined by Western Blot analysis. β-actin was used as a loading control. PLD, pegylated liposomal doxorubicin; PD, progressive disease; IHC, immunohistochemistry.

**Table 1:** *In vivo* responses of GEMM tumours to cisplatin, paclitaxel, pegylated liposomal doxorubicin (PLD), vinorelbine and eribulin. The GEMM tumours were refractory to cisplatin, paclitaxel and PLD as the time to progressive disease (PD) was the same as for vehicle treated mice. PLD and cisplatin failed to demonstrate any meaningful response with no significant difference in median time-to-harvest (TTH) compared to vehicle treatment. Paclitaxel demonstrated modest responses with an increase in median TTH from 15 to 36 days compared to vehicle treated mice (*p* = 0.0101). Improvement in time to PD were seen in tumours treated with vinorelbine (56 days) and eribulin (35 days). This led to a significant improvement of median TTH from 15 days for vehicle treated mice to 81 days with vinorelbine (*p* < 0.0001) and to 46 days with eribulin (*p* < 0.0001). The log-rank test was used for statistical analysis of Kaplan-Meier survival curves (Figure 3a).

| Treatment | Number of mice (n) | Time to progressive disease (PD) (days) | Median time to harvest (TTH) (days) | p value Compared to vehicle | p value Compared to cisplatin | p value Compared to eribulin | p value Compared to paclitaxel | p value Compared to doxorubicin liposomal | p value Compared to vinorelbine | Drug response score |
| --- | --- | --- | --- | --- | --- | --- | --- | --- | --- | --- |
| Vehicle | 25 | 7 | 15 |  |  |  |  |  |  |  |
| Cisplatin | 10 | 7 | 18 | 0.0251 |  | 0.2063 | 0.9378 |  | <0.0001 | Refractory |
| Paclitaxel | 3 | 7 | 36 | 0.0101 | 0.9378 | 0.0067 |  | 0.4855 | 0.0006 | Refractory |
| PLD | 3 | 7 | 29 | 0.0798 | 0.5834 | 0.0043 | 0.4855 |  | 0.0002 | Refractory |
| Vinorelbine | 9 | 56 | 81 | <0.0001 | <0.0001 | 0.0012 | 0.0006 | 0.0002 |  | Responsive |
| Eribulin | 5 | 35 | 46 | <0.0001 | 0.2063 |  | 0.0067 | 0.0043 | 0.0012 | Responsive |

### Eribulin treatment reduced adhesion, invasion and branching of the OCS GEMM cell line

*In vitro* functional assays showed eribulin reduced both adhesion to collagen matrices (Figure 3c; *p*=0.024) and invasion through extracellular matrices of OCS GEMM cells (Figure 3c; *p*=0.0042), compared to DMSO, and reduced branch formation in 3D collagen growth assays (Figure 3d). Western Blot analysis determined a reduction in expression of the mesenchymal markers ZEB1, N-cadherin, vimentin and HMGA2 in OCS GEMM cells exposed to eribulin (Figure 3e).

### A cohort of OCS PDX models with N-MYC/LIN28B pathway up-regulation recapitulated the biphasic and heterogeneous nature of OCS

We next expanded and characterised six PDX models of OCS with varying degrees of carcinoma and sarcoma, all harbouring loss or mutation of TP53 (Figure 4a; Supplementary Table S16). The heterogeneous characteristics of the PDX cohort resembled the human OCS tumour landscape. Furthermore, all PDX models expressed HMGA2, suggesting the N-MYC/LIN28B pathway was up-regulated. Over time, a purely sarcomatous lineage (PH003sarc) arose from the original mixed PH003 model (called PH003mixed). RNAseq data revealed that all PDX had higher HMGA2 expression and EMT scores than the TCGA-OV cohort (Figures 4b and 4c). The most sarcomatous PDX models (PH003sarc and PH592) had higher EMT scores than models containing regions of pure carcinoma (PH419 and PH003mixed). By Western Blot, expression of vimentin was highest in PH142, PH003mixed and PH003sarc. PH419 exhibited the lowest expression of vimentin and PH006 and PH592 had intermediate expression (Figure 4d).

**Figure 4:**
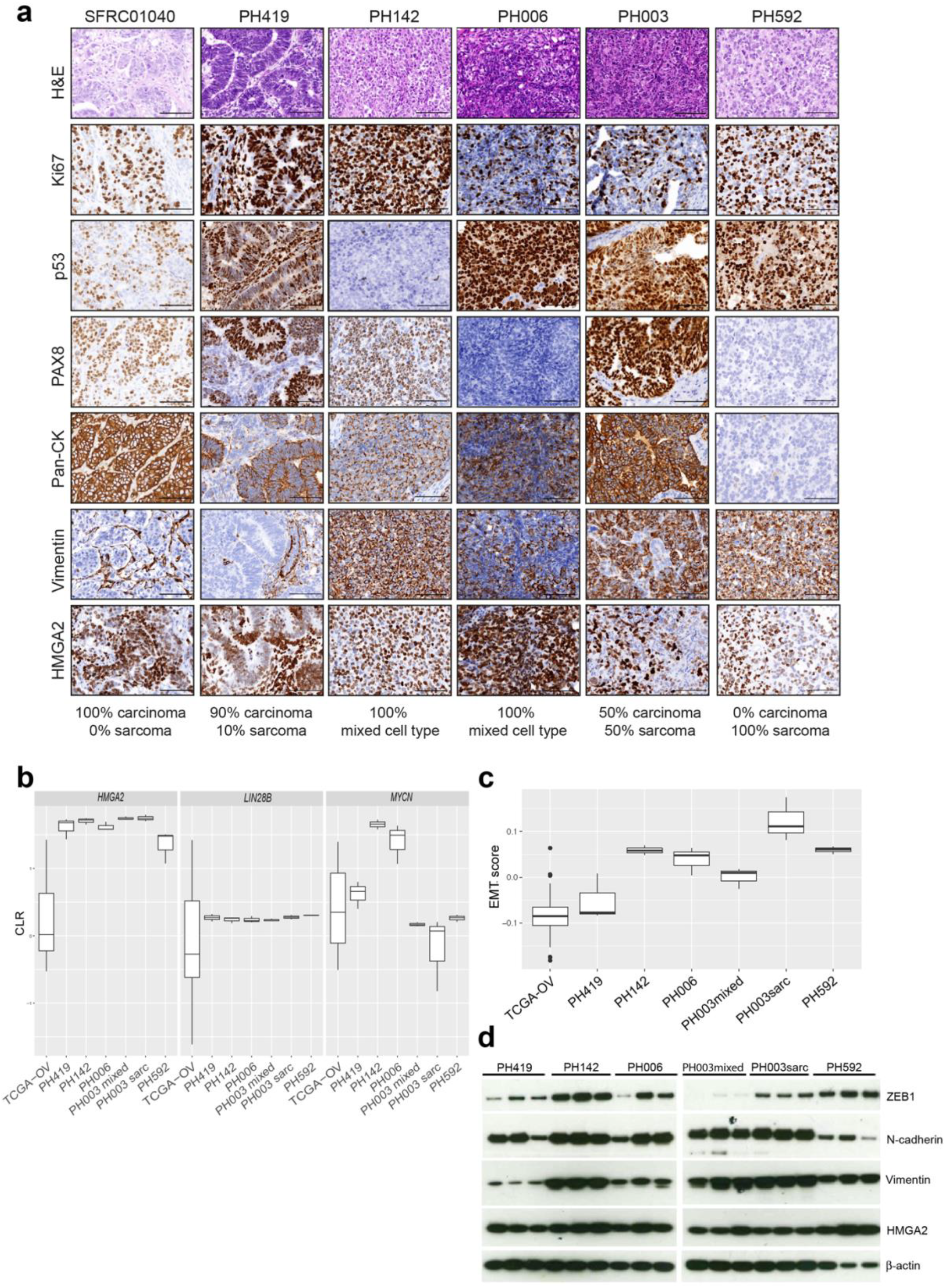
Characterisation of PDX models of OCS with varying proportions of carcinoma and sarcoma. **(A)** Tumours from each PDX model of OCS were assessed by IHC. Representative images of H&E, Ki67, p53, PAX8, Pan-CK, Vimentin and HMGA2 staining are shown. Scale bars represent 100μm. SFRC01040 and PH419 were almost purely carcinoma, PH142, PH006 and PH003 were mixed with both carcinomatous and sarcomatous characteristics (i.e. expressing both Pan-CK and Vimentin) and PH592 was purely sarcomatous, with some epithelial characteristics (i.e. Pan-CK co-expression in some cells). **(B)** Expression of *HMGA2*, *LIN28B* and *MYCN* were determined from RNAseq data for each OCS model (n = 3) compared to ovarian high-grade serous carcinoma samples in TCGA (n = 379). **(C)** EMT scores generated from RNAseq data for tumours from each OCS PDX model are shown compared with EMT scores for ovarian high-grade serous carcinoma samples in TCGA. **(D)** Expression of the mesenchymal markers ZEB1, N-cadherin, Vimentin and HMGA2 in tumours from each OCS PDX model was determined by Western Blot analysis. β-actin was used as a loading control. PDX, patient-derived xenograft; IHC, immunohistochemistry; CK, cytokeratin; TCGA-OV, ovarian high-grade serous carcinomas in TCGA; EMT, epithelial-to-mesenchymal transition; CLR, centred log ratio.

### Platinum based chemotherapy was ineffective in OCS PDX

*In vivo*, four of six PDX were refractory to cisplatin as based on our previously published criteria^59^, failing to achieve any meaningful tumour response and developing progressive disease (PD) whilst on cisplatin therapy (D1-18) (Figure 5a and Supplementary Figure S8). Initial tumour regression was observed in PH142 and SFRC01040 but PD occurred by day 42 and day 60 respectively, defining both as cisplatin resistant PDX^59^ (Table 2).

**Figure 5:**
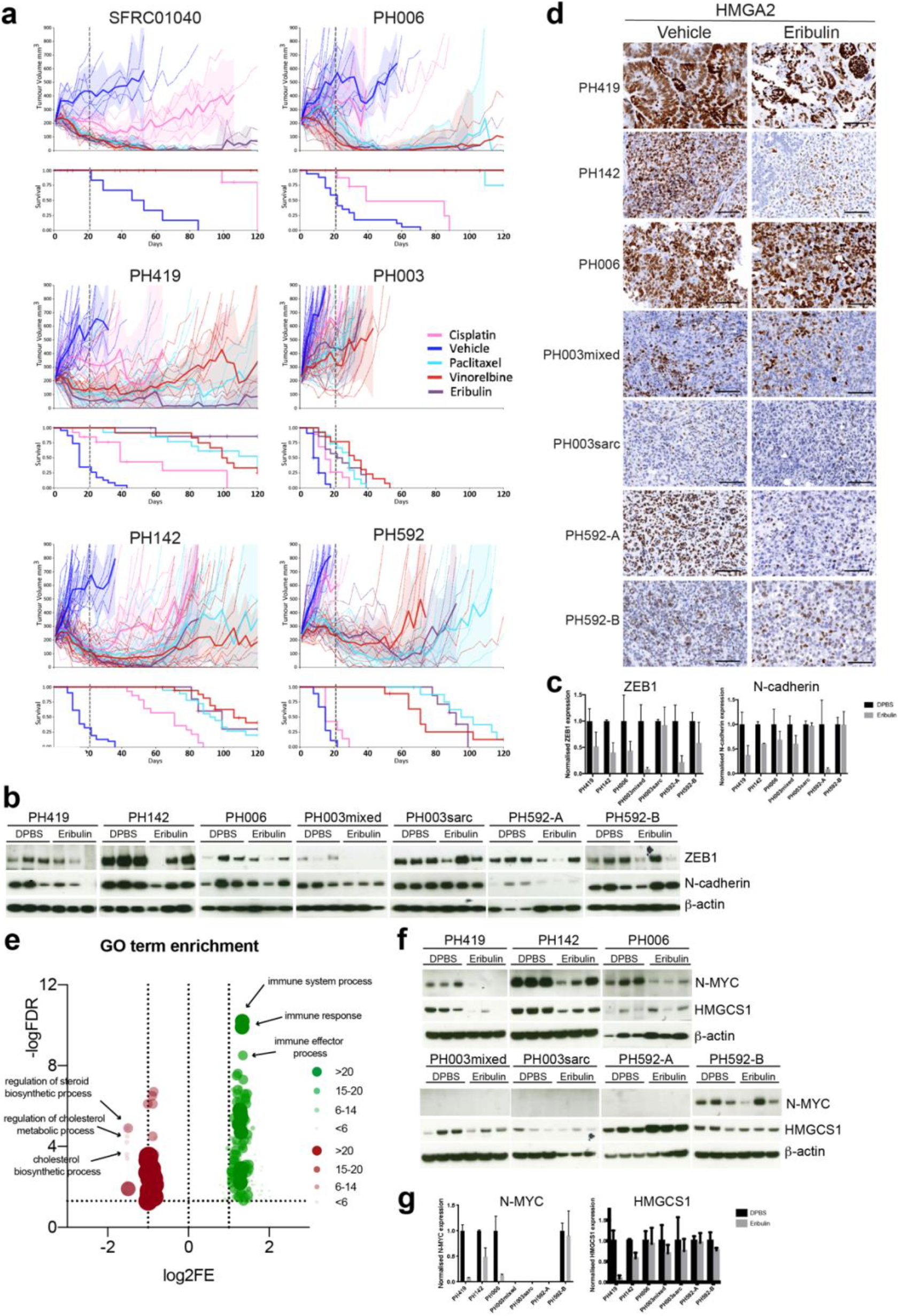
PDX OCS tumours are refractory to cisplatin but display mostly impressive responses to microtubule drugs. **(A)** *In vivo* treatment of OCS PDX tumours with DPBS, cisplatin (4mg/kg), paclitaxel (25mg/kg), vinorelbine (15mg/kg) and eribulin (1.5mg/kg, with the exception of mice harbouring SFRC01040 tumours, which received doses of 1mg/kg). n values for each model are shown in Table 2. Shaded area = 95% confidence interval. More carcinomatous models are shown on the top left and the more sarcomatous models on the bottom right. Time to PD and harvest (TTH) are shown in Table 2. **(B)** Expression of the mesenchymal markers ZEB1 and N-cadherin in tumours from each OCS PDX model after a single dose of vehicle (DPBS) or eribulin was determined by Western Blot analysis. β-actin was used as a loading control. **(C)** Quantification of expression data in (B). **(D)** Expression of HMGA2 in tumours from each OCS PDX model after a single dose of vehicle (DPBS) or eribulin was determined by IHC. Scale bars represent 100μm. **(E)** Analysis of GO terms enriched for down-regulated (red) and up-regulated (green) DEGs. Circle sizes indicate DEGs present in each GO term. DEGs are listed in Supplementary Tables S18 - S21. **(F)** Expression of N-MYC and HMGCS1 in tumours from each OCS PDX model after a single dose of vehicle (DPBS) or eribulin was determined by Western Blot analysis. β-actin was used as a loading control. **(G)** Quantification of expression data in (F). GO, gene ontology; DEG, differentially expressed gene; FDR, false discovery rate.

**Table 2:** *In vivo* responses of OCS PDXs to cisplatin, paclitaxel, vinorelbine and eribulin. Cisplatin failed to achieve any meaningful tumour response in four of six PDX models; PH419, PH006, PH003 and PH592, with a time to development of progressive disease (PD) during cisplatin treatment. PH142 and SFRC01040 demonstrated some response to cisplatin with improvement of median TTH from 15 to 71 days (p < 0.0001) and 53 to 120 days (p = 0.0008), compared to vehicle treated mice, respectively. However, times to PD were less than 100 days (PH142 at 42 days and SRFC01040 at 60 days), therefore these tumours were classified as resistant to cisplatin. Three of six PDX (SFRC01040, PH419 and PH006) were shown to be sensitive to paclitaxel *in vivo*, two PDX (PH142 and PH592) were resistant and one PDX (PH003) was refractory based on the same *in vivo* drug response score as cisplatin. Paclitaxel treated PDX models displayed an impressive improvement in median TTH compared with vehicle treated mice except for PH003 (53 to >120 days for SFRC01040 (*p* = 0.0013), 15 to 120 days for PH419 (*p* < 0.0001), 15 to 95 days for PH142 (*p* < 0.0001), 22 to >120 days for PH006 (*p* < 0.0001), and 15 to 102 days for PH592 (*p* < 0.0001)). Significant improvements in median TTH compared to cisplatin treated mice were observed for four models (39 to 120 days for PH419 (*p* = 0.0036), 71 to 95 days for PH142 (*p* < 0.0001), 39 to >120 days for PH006 (*p* = 0.0024), and 15 to 102 days for PH592 (*p* < 0.0001)). Three of six OCS PDX (SFRC01040, PH142 and PH006) were sensitive, two PDXs (PH419 and PH592) were resistant and one PDX (PH003) was refractory to vinorelbine treatment. Significant improvements of median TTH compared with vehicle treated mice were observed for all models treated with vinorelbine except for PH003 (53 to >120 days for SFRC01040 (*p* = 0.0033), 15 to 99 days for PH419 (*p* < 0.0001), 15 to 106 days for PH142 (*p* < 0.0001), 22 to >120 days for PH006 (*p* < 0.0001), and 15 to 71 days for PH592 (*p* < 0.0001)). There were significant improvements in median TTH compared with cisplatin treated mice were also observed for four models (120 to >120 days for SFRC01040 (*p* = 0.1232), 39 to 99 days for PH419 (*p* = 0.0151), 71 to 106 days for PH142 (*p* < 0.0001), 39 to >120 days for PH006 (*p* = 0.0012), and 15 to 71 days for PH592 (*p* < 0.0001)). Three of six OCS PDX models (SFRC01040, PH419 and PH006) were sensitive, two PDX (PH412 and PH592) were resistant and one PDX (PH003) was refractory to eribulin treatment. Significant improvements of median TTH were observed in eribulin treated mice compared to vehicle for five models (53 to >120 days for SFRC01040 (*p* = 0.0015), 15 to 99 days for PH142 (*p* < 0.0001), 22 to >120 days for PH006 (*p* = 0.0005), 15 to >120 days for PH419 (*p* < 0.0001), and 15 to 92 days for PH592 (*p* = <0.0001)). Lastly, significant improvements in median TTH compared with cisplatin treated mice were also observed for four models (120 to >120 days for SFRC01040 (p = 0.0511), 39 to >120 days for PH419 (*p* = 0.0016), 71 to 99 days for PH142 (*p* = 0.0042), 39 to >120 days for PH006 (*p* = 0.0079), 15 to 25 days for PH003 (*p* = 0.0044), and 15 to 92 days for PH592 (*p* < 0.0001)). The log-rank test was used for statistical analysis of Kaplan-Meier survival curves (Figure 5a).

| PDX model | Treatment | Number of mice (n) | Time to progressive disease (PD) (days) | Median time to harvest (TTH) (days) | p value Compared to vehicle | p value Compared to cisplatin | p value Compared to eribulin | p value Compared to paclitaxel | p value Compared to vinorelbine | Drug response score (Topp et al) |
| --- | --- | --- | --- | --- | --- | --- | --- | --- | --- | --- |
| SFRC01040 | Vehicle | 8 | 7 | 53 |  |  |  |  |  |  |
|  | Cisplatin | 8 | 60 | 120 | 0.0008 |  |  |  |  | Resistant |
|  | Eribulin | 7 | >120 | >120 | 0.0015 | 0.0511 |  | >0.9999 | >0.9999 | Sensitive |
|  | Paclitaxel | 7 | >120 | >120 | 0.0013 | 0.0511 | >0.9999 |  | >0.9999 | Sensitive |
|  | Vinorelbine | 6 | >120 | >120 | 0.0033 | 0.1232 | >0.9999 | >0.9999 |  | Sensitive |
| PH419 | Vehicle | 23 | 7 | 15 |  |  |  |  |  |  |
|  | Cisplatin | 13 | 7 | 39 | <0.0001 |  |  |  |  | Refractory |
|  | Eribulin | 8 | >120 | >120 | <0.0001 | 0.0016 |  | 0.0875 | 0.0446 | Sensitive |
|  | Paclitaxel | 14 | 112 | 120 | <0.0001 | 0.0036 | 0.0875 |  | 0.6489 | Sensitive |
|  | Vinorelbine | 12 | 80 | 99 | <0.0001 | 0.0151 | 0.0446 | 0.6489 |  | Resistant |
| PH142 | Vehicle | 31 | 7 | 15 |  |  |  |  |  |  |
|  | Cisplatin | 19 | 42 | 71 | <0.0001 |  |  |  |  | Resistant |
|  | Eribulin | 10 | 77 | 99 | <0.0001 | 0.0042 |  | 0.8113 | 0.4092 | Resistant |
|  | Paclitaxel | 22 | 57 | 95 | <0.0001 | <0.0001 | 0.8113 |  | 0.1134 | Resistant |
|  | Vinorelbine | 19 | 120 | 106 | <0.0001 | <0.0001 | 0.4092 | 0.1134 |  | Sensitive |
| PH006 | Vehicle | 17 | 7 | 22 |  |  |  |  |  |  |
|  | Cisplatin | 9 | 7 | 39 | 0.0064 |  |  |  |  | Refractory |
|  | Eribulin | 6 | >120 | >120 | 0.0005 | 0.0079 |  | 0.4795 | >0.9999 | Sensitive |
|  | Paclitaxel | 7 | >120 | >120 | <0.0001 | 0.0024 | 0.4795 |  | 0.3173 | Sensitive |
|  | Vinorelbine | 7 | >120 | >120 | <0.0001 | 0.0012 | >0.9999 | 0.3173 |  | Sensitive |
| PH003 | Vehicle | 23 | 7 | 8 |  |  |  |  |  |  |
|  | Cisplatin | 19 | 7 | 15 | 0.0005 |  |  |  |  | Refractory |
|  | Eribulin | 14 | 7 | 25 | 0.0003 | 0.0044 |  | 0.9612 | 0.1339 | Refractory |
|  | Paclitaxel | 16 | 7 | 29 | <0.0001 | 0.0025 | 0.9612 |  | 0.0666 | Refractory |
|  | Vinorelbine | 13 | 18 | 32 | <0.0001 | 0.0005 | 0.1339 | 0.0666 |  | Refractory |
| PH592 | Vehicle | 18 | 7 | 15 |  |  |  |  |  |  |
|  | Cisplatin | 7 | 7 | 15 | 0.0335 |  |  |  |  | Refractory |
|  | Eribulin | 8 | 80 | 92 | <0.0001 | <0.0001 |  | 0.2838 | 0.3070 | Resistant |
|  | Paclitaxel | 8 | 88 | 102 | <0.0001 | <0.0001 | 0.2838 |  | 0.2183 | Resistant |
|  | Vinorelbine | 9 | 63 | 71 | <0.0001 | <0.0001 | 0.3070 | 0.2183 |  | Resistant |

### Microtubule-targeting agents, such as paclitaxel, vinorelbine and eribulin, were effective in OCS

Microtubule-targeting agents induced tumour regression and showed an improvement of median TTH in most OCS PDX models. Three PDX (SFRC01040, PH419 and PH006) were classified as sensitive to paclitaxel according to the same criteria used for cisplatin^59^, two were resistant (PH142 and PH592) and one was refractory (PH003) (Figure 5a and Supplementary Figure S8). Indeed, all models displayed an improvement in median TTH compared with vehicle, except for PH003, with four models also displaying an improvement in median TTH compared with cisplatin (Table 2).

Three of six OCS PDX (SFRC01040, PH142 and PH006) were sensitive to vinorelbine, two resistant (PH419 and PH592) and one refractory (PH003) (Figure 5a and Supplementary Figure S8). Interestingly, the more sarcomatous PDX models, PH003 and PH592, were less sensitive to vinorelbine than were the more carcinomatous models. Significant improvements in median TTH compared with vehicle were observed for all models except PH003, and in four models compared with cisplatin (Table 2).

Lastly, three of six PDX models (SFRC01040, PH419 and PH006) were sensitive to eribulin treatment, two were resistant (PH142 and PH592) and one was refractory (PH003) (Figure 5a and Supplementary Figure S8). Interestingly, near complete responses to eribulin were observed in three PDX (SFRC01040, PH419 and PH006). Significant improvements in median TTH compared to vehicle and cisplatin were observed for five models and four models, respectively (Table 2). Eribulin treatment of PH592, which was predominantly sarcomatous, resulted in significant tumour stabilisation to 40 days followed by marked tumour regression between days 60 to 80 before rapid disease progression. Even for the most aggressive model, PH003, eribulin treatment resulted in a statistically significant improvement in median TTH, albeit of short duration (8 days (vehicle) vs 25 days (eribulin) (*p*=0.0003) and 15 days (cisplatin) vs 25 days (eribulin) (*p*=0.0044)) (Table 2).

It was noted that one lineage of the sarcomatous PDX PH592 was markedly more sensitive to cisplatin treatment, denoted PH592-B, compared with the more cisplatin resistant, PH592-A lineage (median TTH of 15 days (PH592-A) vs 71 days (PH592-B); *p*<0.0001) and similarly, was more sensitive to eribulin (92 days (PH592-A) vs 102 days (PH592-B); *p*=0.0240) (Supplementary Figure S9 and Supplementary Table S17).

### *In vivo* eribulin treatment reduced the expression of mesenchymal markers, including HMGA2, in OCS PDX tumours

PDX tumours were harvested one week after mice received a single dose of eribulin (or DPBS vehicle control) and expression of EMT markers was assessed by western blotting and IHC. Eribulin reduced expression of the mesenchymal marker HMGA2 as well as ZEB1 and N-cadherin in most models (Figures 5b-d; Supplementary Figure S10).

### Reduced cholesterol biosynthesis and increased immune activation was implicated in the response of a subset of OCS to eribulin

RNAseq analysis of PDX tumours harvested one week after a single dose of eribulin (Supplementary Table S15) indicated significant down-regulation of genes related to the GO terms protein targeting to membrane, translational initiation, and regulation of cholesterol biosynthesis, and up-regulation of genes related to the GO term immune activation (Figure 5e; Supplementary Tables S18-S21). Interestingly, significantly down-regulated genes included eight genes involved in the mevalonate (MVA) pathway, which plays a key role in cholesterol biosynthesis: *SREBF2*, *HMGCR*, *HMGCS1*, *MVK*, *LDLR*, *INSIG1*, *IDI1*, *FDFT1*. Expression of hydroxymethylglutaryl-CoA synthase (HMGCS1), a key enzyme in the MVA pathway, was assessed by Western Blot analysis and found to be reduced in PH419, PH142 and PH592-B following a single dose of eribulin (Figures 5f and 5g). In neuroblastoma, which is commonly driven by *MYCN* amplification, there is also increased activation of the MVA pathway and apparent reliance on this pathway for survival^71^. We hypothesise that N-MYC is a key driver of OCS, implicating the MVA pathway in OCS cell survival and drug resistance. Notably, in three of the PDX models where there was no change in HMGCS1 expression, PH003mixed, PH003sarc and PH592-A, RNAseq showed these models had low expression of *MYCN* with maintained levels of *LIN28B* (Figure 4b), and expression of N-MYC was almost undetectable by western blot (Figures 5f and 5g). These tumours also had the poorest relative response to eribulin *in vivo* (Figure 5b).

## Discussion

OCS is a rare, heterogeneous and clinically aggressive cancer, with poorer overall survival than HGSC despite a similar mutation and copy number profile^60,61.^ Nearly all patients with metastatic OCS, despite initial response to standard-of-care platinum-based chemotherapy, will succumb to their cancer due to early relapse of disease^62^. There is no effective second-line therapy available owing to its multi-drug resistant behaviour^62^. Additionally, the biphasic nature of OCS and a poor understanding of how these tumours develop has hindered progress in the development of effective treatment options.

There have been few previous molecular studies of OCS tumours where the carcinomatous and sarcomatous components have been micro-or macro-dissected^4,5,7.^ Two recent studies performed whole exome sequencing on separated components of OCS tumours, but on no more than four tumours each^8,9.^ Here, we analysed 377 genes (for mutations, copy number, or both) in 18 OCS tumours where the carcinomatous and sarcomatous components were analysed independently along with associated metastases, where available. We found mutations commonly identified in OCS, with the initial or truncal mutation likely to occur in *TP53*. In nearly all of the cases, the same *TP53* mutation was identified in all sites available; carcinoma, sarcoma and metastasis. Independent sites then developed additional mutations in most cases. Through this we definitively determined that OCS tumours in our cohort were monoclonal. Furthermore, we carried out RNAseq analysis, which has not previously been achieved for the independent components in OCS. The carcinomatous component was found to have a significantly higher EMT score than conventional HGSC, indicating these tumours may have been primed to undergo sarcomatous transformation early in carcinogenesis. Together, these data support the conversion theory for OCS histogenesis and highlight the basis of the aggressive behaviour of tumours that look, at a genomic level, indistinguishable from routine HGSC but behave like the very worst prognostic outliers. Future studies, utilising high-resolution single cell sequencing approaches, are required to prove this definitively. Nevertheless, this study, in keeping with existing published evidence^8,9^, further emphasises the potential key role of EMT in OCS tumorigenesis and biological behaviour. This study also highlights the potential downfall of treating women with OCS in the same way as HGSC (with the exception of *BRCA1/2*-mutant OCS cases for whom PARP inhibitor therapy is reasonable^63^), as we have shown that despite the genomic similarity, OCS are phenotypically distinct, particularly with regard to drug responses and mesenchymal characteristics.

Similarities between OCS and the C5 molecular subtype of HGSC, as defined by Tothill *et al*^38^, include the poor clinical outcome and link to drug resistance, as well as deregulation of the *let-7* pathway^36^, which we have called the N-MYC/LIN28B pathway. We confirmed that *LIN28B* and *HMGA2* were significantly up-regulated in our cohort of 18 OCS tumours compared to HGSC. This suggested that the N-MYC pathway is important in the development and maintenance of OCS. Using this knowledge, we developed a GEMM of OCS by overexpressing *Lin28b* and inhibiting p53 in PAX8^+^ FTSECs. While the OCS GEMM tumours exhibited high expression of *Lin28b* and *Mycn*, the derived cell line displayed high expression of *Lin28b* and *Hmga2*, indicating that we had generated two closely related pre-clinical models of OCS with different characteristics. This demonstrates the complexity of the N-MYC pathway, as was also indicated by the RNAseq data from our patient samples. Observed expression of this pathway depends on multiple feedback loops and influences from outside the pathway, such as transcription factors, and occurs at the level of transcription and translation, frequently resulting in complex relationships^64^.

These models were used to compare the current standard-of-care treatments for OCS with novel treatments, including the unique microtubule-targeting drug, eribulin, that has been shown to reverse EMT^42–46^, and has demonstrated improved efficacy against metastatic breast cancer, soft-tissue sarcoma and ovarian cancer^48,65,66.^ While the GEMM tumours were refractory to cisplatin, paclitaxel and PLD *in vivo*, they were responsive to vinorelbine and eribulin. Disease stabilisation was achieved with both vinorelbine and eribulin, suggesting a longer progression free survival may be achieved with these drugs in patients. Furthermore, after just a single dose of eribulin, a notable decrease in tumour cell proliferation was observed. To test the mechanism of action of eribulin, we used the GEMM cell line and observed significantly reduced adhesion and invasion following eribulin treatment, which corresponded with an inability of these cells to branch out in 3D matrix. Finally, an impressive reduction in expression of the mesenchymal markers ZEB1, N-cadherin, and Vimentin was observed in cells exposed to eribulin while growing on collagen, as well as in HMGA2. These results are consistent with a previous report where eribulin reversed the process of EMT, thus reducing the mesenchymal characteristics of breast cancer cells^44^.

A cohort of molecularly annotated OCS PDX models, closely resembling human OCS, were characterised for novel drug efficacy. We obtained six PDX models of OCS with a range of carcinoma and sarcoma characteristics. RNAseq analysis of these tumours indicated all OCS had higher EMT scores than HGSC, with the most carcinomatous model PH419 having the lowest EMT score and the most sarcomatous model PH003sarc having the highest EMT score. At the protein level, PH003sarc also had the highest expression of the mesenchymal markers N-cadherin and Vimentin. Interestingly, the two models containing mixed cells, PH142 and PH006 also had high expression of N-cadherin, Vimentin and ZEB1. This matched their high EMT scores obtained from the RNAseq data and indicated that pathology alone was insufficient to determine the level of sarcomatous transformation occurring in each OCS model.

Anti-microtubule agents, as a class of drug, were more effective than platinum-based chemotherapy in our diverse cohort of OCS PDX models. Interestingly, the proportion of carcinoma and sarcoma did not appear to correlate with anti-microtubule drug sensitivity. Whereas for cisplatin, the more carcinomatous PDX had some initial response, while the most sarcomatous PDX were completely refractory. Indeed, impressive responses were observed for almost all PDX to the microtubule-targeting drugs, paclitaxel, vinorelbine and eribulin. PDX PH003 was the one exception where tumours remained refractory to all treatment regimens tested. This drug-refractory PDX was later found to lack N-MYC expression, representing a particularly aggressive subtype of OCS, corresponding to rapidly progressive disease in the patient^67^. Possibly as a consequence of lacking N-MYC, PH003 tumours also exhibited the lowest expression of HMGA2. Interestingly, PH952-A, the more drug-resistant lineage of PH592, also lacked expression of N-MYC, whereas it was expressed in the more drug sensitive lineage, PH592-B. Eribulin is known to reverse EMT characteristics, and indeed we observed a decrease in N-cadherin and ZEB1 protein expression in most models following a single dose of eribulin. Reduced ZEB1 and N-cadherin expression was not displayed in all of our models by IHC, which could be explained by the region of the tumour analysed. Importantly, after a single dose of eribulin, a decrease in the expression of HMGA2 was observed in PH419, PH142, PH003sarc, PH592-A and PH592-B tumours. We hypothesised that eribulin interferes with the N-MYC pathway, leading to a reduction in the mesenchymal characteristics of OCS tumours, including down-regulation of HMGA2.

To better understand the mechanism of action of eribulin in our PDX models, we carried out RNAseq analysis after a single dose of eribulin and found a significant reduction in the expression of genes involved in the MVA pathway and a significant up-regulation of genes involved in activation of immune responses. Cholesterol synthesis is important for cell membrane biogenesis and, therefore, cancer cell growth and proliferation^68^. Furthermore, there are indications that cholesterol is involved in EMT^69,70^. We hypothesised that the mechanism by which eribulin reduces EMT characteristics was by inhibiting cholesterol synthesis. To substantiate this finding, we analysed the expression of a key enzyme in the MVA pathway, HMGCS1, in PDX after a single dose of eribulin. We saw a reduction of HMGCS1 expression in four PDX. However, the three PDX that did not display reduced HMGCS1 expression after eribulin treatment lacked expression of N-MYC. We hypothesise that, as in neuroblastoma^71^, N-MYC drives OCS cell survival and drug resistance through the MVA pathway. This pathway appears to be targeted, at least in part, by eribulin, leading to reduced expression of N-MYC, HMGA2, and reversal of EMT characteristics.

The involvement of the MVA pathway in OCS survival suggests that statins may have therapeutic potential. However, in future studies of statins in OCS, it would be important to consider the tightly controlled SREBP2-mediated feedback loop, which acts to increase the expression of MVA pathway genes^72^. This is a potential mechanism of drug resistance, indeed it has been implicated in cisplatin resistance in ovarian cancer^73^, which may be overcome by combination regimens^74^. Considering, as we have demonstrated here, that eribulin can reduce the expression of many genes in the MVA pathway, combining eribulin with statins could potentially overcome resistance that might arise with statin therapy alone.

High levels of cholesterol have also been shown to play a protective role in cancer cells through inhibiting immune surveillance^75^. Indeed, in our OCS PDX models we also observed a significant increase in the expression of genes involved in immune activation following eribulin treatment. Thus, eribulin may initiate anti-tumour immune responses in OCS, as has been observed in other tumour types^45,47,76.^ Therefore, early phase clinical trials in OCS for eribulin as a single agent and in combination with immunotherapy should be initiated to improve treatment options for OCS.

## Patients and Methods

### Study conduct, survival analyses and patient samples

Overall survival was calculated from the date of diagnosis to the date of death or the last known clinical assessment. Overall survival was calculated by log-rank test (Mantel-Cox) using Prism v8.0 (GraphPad, San Diego, CA).

Formalin-fixed paraffin-embedded (FFPE) specimens were identified from the pathology archives of Queen Elizabeth University Hospital, Glasgow, UK. Following review by an expert gynaecological pathologist, areas of carcinoma and sarcoma were marked for macro-dissection.

### Panel Sequencing

Libraries for sequencing were prepared from genomic DNA (gDNA) obtained from 5 × 10μm macro-dissected FFPE sections. A total input of 50-200ng per sample was used based on quantification with a Quant-iT PicoGreen dsDNA Assay Kit (Invitrogen, Carlsbad, CA, USA). Each DNA sample was sheared using a Covaris LE220 focused-ultrasonicator (Covaris, Woburn, MA, USA) with the following settings: PIP450, Cycles/Burst 200, Duty Factor 15%, Water Level of 6, shearing time of 400 seconds (executed as 350 seconds, followed by a further 50 seconds using the same settings). Pre-capture sample libraries were prepared on the SciClone G3 NGS Workstation (Perkin Elmer, Waltham, MA, USA) using SureSelect XT standard automated protocol (Agilent Technologies, Santa Clara, CA, USA) for 200ng samples. Pre-capture sample libraries were quantified with the Quant-iT PicoGreen dsDNA Assay Kit. Quantification data were used to normalise all sample libraries to 750ng in a total volume of 26.4μl; a full 26.4μl of sample library was brought forward for libraries with a total concentration too low to make this possible. Normalised pre-capture sample libraries were then captured using 120nt biotinylated custom RNA baits from a proprietary SureSelect XT custom 6-11.9Mb panel (Agilent Technologies, Santa Clara, CA, USA). Captured libraries were processed as a large panel, since more than 3Mb of sequence was intended for capture, and were incubated overnight to facilitate hybridisation, as per manufacturer’s protocol. Captured sample library sequences were extracted from solution, cleaned up and prepared for post-capture PCR. Post-capture PCR incorporated primers with unique 8-bp indexes (Agilent Technologies, Santa Clara, CA, USA) for multiplexing. Amplified capture libraries were cleaned up on a Zephyr G3 NGS Workstation (Perkin Elmer, Waltham, MA, USA), using a post-PCR SPRI bead clean-up protocol (Agilent Technologies, Santa Clara, CA, USA), to produce final capture libraries. Final captured-libraries were quantified with the Quant-iT PicoGreen dsDNA Assay Kit and assessed for size distribution and quality on a LabChip GX DNA High Sensitivity Chip (Perkin Elmer, Waltham, MA, USA). 8 uniquely indexed sample libraries were pooled per lane of a HiSeq 4000 flow cell. Pools were clustered to the flow cell using a cbot 2 system and sequenced on a HiSeq 4000 (Illumina, San Diego, CA, USA) as per manufacturer’s instructions to generate 2×75bp reads.

### Panel design and analysis

Genes for inclusion in the custom panel were selected from publicly available databases (including the Cancer Gene Census (CGC)^77^, Database of Curated Mutations (DoCM)^78^ and Vogelstein et al’s analysis of COSMIC^79^) as well as unbiased statistical screens^80–83^. For genes where driver events are mainly substitutions (e.g. *MAP2K1, GNA11, MTOR, NRAS*), the coding exons were included in the panel design. For genes where driver events are mainly copy number alterations (e.g. *CCND2, CCNE1, FGF3, MDM2*), approximately 20 marker SNPs spanning the gene footprint were included in the panel design. For key tumour suppressor genes (e.g. *BRCA1, BRCA2, CDKN2A, NF1, PTEN, RB1*) where driver events could be any inactivating sequence-level, structural or copy number change, the entire gene footprint was included in the panel design. In total, this panel assays 217 genes for coding sequence mutations, 137 genes for copy number state, and 23 genes for all genomic events. In addition, SNPs spaced approximately 1Mb apart throughout the genome were included to give a genome-wide copy number profile. Total sequence capture size was 3.465MB.

Sequencing data were analysed using HOLMES, a proprietary pipeline that uses a Snakemake^84^ workflow to run the following data processing steps: 1) bcl2fastq v2.19.1 (https://support.illumina.com/sequencing/sequencing_software/bcl2fastq-conversion-software.html) or fastq generation and adapter trimming. 2) bwa mem v0.7.15^85^ for alignment to GRCh38 and biobambam v2.0.72^86^ for sorting, indexing, duplicate marking and duplicate removal. 3) samtools stats v1.5^87^ to generate QC metrics. 4) Shearwater/deepSNV v1.22.0^88,89^ to call point mutations from properly paired reads only, using all samples from this project as the cohort, with the following filters then applied: there must be no evidence for the same mutation in the matched normal, the average mapping quality of reads supporting the variant must be ≥ 20, the variant must not be present in the 1000 Genomes Project, at least a third of bases reporting the mutation must have a base quality of at ≥20, the allele frequency of the mutation must be at least 5%, there must be ≥ 20 reads covering the variant position and at least 3 must contain the mutation, the ratio of forward to reverse reads containing the mutation must be between 0.15 and 6.67 inclusive, not more than 10% of reads containing the mutation can contain an indel with 10 bp of the variant position. 5) Pindel v0.2.5b8^90^ to call indels, with the following filters applied: only reads with a mapping quality of ≥10 are used as anchors, the variant must not be present in the 1000 Genomes Project, there must be no evidence for the same indel in the matched normal if it is >4bp long or the allele frequency of the indel must be 10x higher in the tumour than in the matched normal if it is ≤4bp long, at least 3 reads must report the indel, the same indel must not be called in any of the matched normals in this project, the allele frequency of the indel must be at least 5%, the ratio of forward to reverse reads containing the indel must be between 0.1 and 10 inclusive. 6) Annotation of both substitutions and indels with CAVA v1.2.2.^91^. 7) GeneCN v1.0 as described previously^92^ for calling the copy number state of genes and generate genome-wide copy number plots. Samples with high levels of noise, identified as those with large standard deviations within each genomic feature, were excluded from copy number analysis. This excluded WW00153c, WW00169a and WW00170c from individual analyses. 8) Brass (brass-groups command only) v5.3.3 (https://github.com/cancerit/BRASS) for calling structural variants (SVs) with the following filters applied: only reads with a mapping quality of ≥ 10 are considered to support an SV, ≥ 10 read pairs must support an SV, SVs must not fall within the mitochondrial genome or any unplaced or alternative contig, there must be no evidence for the same SV in the matched normal, there must be no evidence of the same SV in any of the matched normals in this project.

To compare the copy number profiles of the sarcoma and carcinoma components, GeneCN was modified to use R’s scale function to centre and scale the data to account for different cellularity between samples. One profile was then subtracted from the other and calling performed on the resulting difference between the two profiles.

### RNA sequencing library generation and sequencing

RNA-seq libraries for the FFPE OCS patient cohort were generated as described in TruSeq Stranded Total RNA Sample Preparation Guide (Illumina, part no. 15031048 Rev. E October 2013) using Illumina TruSeq Stranded Total RNA LT sample preparation kit. Ribosomal depletion step was performed on 500ng of total RNA using Ribo-Zero Gold (Illumina, 20020598 and 20020492). Heat fragmentation step was adjusted depending on RIN score (0 to 8 min) aimed at producing libraries with an insert size between 120-200bp. First strand cDNA was synthesised from the enriched and fragmented RNA using SuperScript II Reverse Transcriptase (Thermofisher, 18064014) and random primers. Second strand synthesis was performed in the presence of dUTP. Following 3’ adenylation and ligation of adaptors to the dsDNA, libraries were subjected to 13 cycles of PCR. RNA-seq libraries were quantified using PicoGreen assay (Thermofisher, P11496) and sized and qualified using an Agilent 4200 TapeStation with Agilent D1000/High sensitivity ScreenTape (Agilent, 5067-5584). Libraries were normalised to 4nM and pooled before clustering using a cBot2 followed by 75bp paired-end sequencing on a HiSeq 4000 sequencer (Illumina).

RNAseq_V2 processed counts for HGSC from TCGA (TCGA-OV cohort) (n=396) were downloaded from the GDC portal (https://portal.gdc.cancer.gov/), version available on 3^rd^ June 2019. In total, there were n = 374 files for primary tumours and n = 5 recurrent tumours. Counts were normalised across samples using DESeq2’s median of ratios method^93^. Carcinosarcoma RNAseq data (n = 27) underwent QC and was found to be satisfactory as per the parameters in FastQC (v.0.11.8 available at http://www.bioinformatics.babraham.ac.uk/projects/fastqc/). Using quasi-mapping method in Salmon version 0.8.2^94^, RNAseq data was aligned to GRCh37 Ensembl release 75^95^ transcriptome. Only those samples where rRNA reads account for less than 20% of the total reads were retained for the downstream analyses, n = 22 (n = 5 samples were excluded, Supplementary Figure S3). Differentially expressed genes (DEGs) between the carcinoma and sarcoma components were derived using the DESeq2 package^93^. The Database for Annotation, Visualization and Integrated Discovery (DAVID) online Functional Annotation Tool was used for functional annotation of Differentially Expressed Genes (DEG).

For EMT gene set enrichment analysis, SingScore^96^ was used with a representative directional gene set^49^. Counts were normalised by rank normalisation^98^ followed by the centred log-ratio transformation^97^. All analyses, statistical tests, and plots were generated in R version 3.3.3 unless specified otherwise.

RNAseq libraries for the PDX tumours were prepared using TruSeq RNA Library Prep Kit v2 (Illumina), and the sequencing was performed on the Novaseq platform to read length of 100 bp (Australian Genome Research Facility). Reads were mapped to the GRCh38 Ensembl release 97 transcriptome and quantified using Kallisto^98^. Counts were normalised and EMT gene set enrichment analysis undertaken as above. DEGs between treated and untreated samples were derived using matching methods across batch and model to correct for batch effects and inherant model differences. *p*-values for DEGs were computed under a normality assumption. Topconfects^99^ was used to calculate lower bounds on the effect sizes with 95% confidence.

### Generation of a genetically-engineered mouse model (GEMM)

The *Pax8-rtTA* strain (C57BL/6 background) was a kind gift from Prof Ronny Drapkin (University of Pennsylvania, Department of Obstetrics and Gynecology, US). The *kai-tetOCre* strain (FVB background) was a kind gift from Prof Jane Visvader (WEHI, Melbourne, Australia) originally sourced from the Osaka Bioscience Institute, Japan. The *LSL-Lin28b* strain (mixed 129X1/SvJ background) was a kind gift from Prof Johannes H. Schulte (University Hospital Essen, Germany; Supplementary Table 14). Mice with multiple transgenes were generated through crossing and breeding mice on a mixed background, predominantly FVB/NJ and C57BL/6. Genotyping was performed using custom designed probes (TransnetYX, Inc; Supplementary Table S15). Activation of the transgenes was achieved through the administration of doxycycline, either by chow (Glen Forrest Stockfeeders SF08-026) or through drinking water (Sigma-Aldrich) at 600mg/kg or 0.2mg/ml respectively. Mice age between 3 weeks and 7 weeks and were treated for 2 weeks to allow adequate doxycycline exposure. Fallopian tubes were carefully micro-dissected, gently minced, and transplanted into the ovarian bursae of CBA/nu mice.

### Immunohistochemistry

Formalin fixed tumour samples were sectioned stained with haematoxylin and eosin (H&E) as well as being sent for automatic immunostaining using the Ventana BenchMark Ultra fully automated staining instrument (Roche Diagnostics, USA). The following antibodies were used: anti-Ki67 (mouse: D3B5, Cell Signalling; human: MIB-1, Dako), anti-PAX8 (polyclonal, Proteintech), anti-p53 (mouse: CM5, Novacastra; human: DO-7, Dako), anti-PanCK (mouse: polyclonal, Abcam; human: AE1/3, Dako), anti-Vimentin (D21H3, Cell Signalling), anti-HMGA2 (D1A7, Cell Signalling), anti-N-cadherin (polyclonal, Abcam), and anti-ZEB1 (polyclonal, NovusBio). H&E and IHC slides were scanned digitally at 20x magnification using the Pannoramic 1000 scanner (3DHISTECH Ltd.). High definition images were uploaded into CaseCenter (3DHISTECH Ltd.) and images were processed using FIJI image application^100^.

### Western Blot Analysis

Tumours homogenised in ice-cold RIPA buffer (50 mM Tris; pH7.5, 150 mM NaCl, 1% NP40, 0.5% sodium deoxycholate, 0.1% SDS in H_2_O, supplemented with a complete mini protease inhibitor cocktail tablet (Roche)) using Precellys Ceramic Kit tubes in the Precellys 24 homogenising instrument (Thermo Fisher Scientific). Proteins from lysates were separated on NuPAGE^®^ Novex^®^ Bis-Tris 10% gels (Invitrogen). Gels were transferred onto PVDF membranes using the iBlot™ Transfer system (Thermo Fish Scientific). Membranes were probed with antibodies specific for ZEB1, N-cadherin, Vimentin, HMGA2 (all as mentioned previously), N-MYC (D1V2A, Cell Signalling), HMGCS1 (A-6, Santa Cruz), or β-actin (AC-15, Sigma).

### Sample processing for RNA and DNA

Total RNA was isolated from snap-frozen cells or tumours using the Direct-zol™ RNA Miniprep kit (Zymo Research) as per manufacturer’s instructions. Tumour DNA was isolated from snap-frozen cells or tumours using the QIAamp DNA mini kit (Qiagen) as per manufacturer’s instructions.

### *In vivo* studies

PDX SFRC01040 was obtained from the Royal Women’s Hospital under the Australian Ovarian Cancer Study and generated by mixing tumour cells isolated from ascites with Matrigel Matrix (Corning) and transplanting subcutaneously into NOD/SCID/IL2Rγnull recipient mice (T1 = passage 1). All other PDXs were rescued through transplanting fragments of cryopreserved tumour tissue subcutaneously from PDXs generated in the Mayo Clinic (USA). GEMM tumours were generated as described above. Recipient mice bearing T2-T7 PDX or GEMM tumours (180-300 mm^3^ in size) were randomly assigned to cisplatin (Pfizer), pegylated liposomal doxorubicin (PLD; Janssen-Cilag Pty. Ltd.), paclitaxel (Bristol-Myers Squibb), vinorelbine (Pfizer), eribulin (Eisai Co., Ltd.), or vehicle treatment groups. *In vivo* cisplatin treatments were performed by intraperitoneal (IP) injection of 4 mg/kg given on days 1, 8 and 18. The regimen for PLD treatment was by IP injection once a week for three weeks at 1.5 mg/kg. The regimen for paclitaxel treatment was by IP injection twice a week for three weeks at 25 mg/kg. The regimen for vinorelbine was by intravenous injection of 15 mg/kg at days 1, 8 and 18. The regimen for eribulin treatment was by IP injection three times a week for three weeks at 1.5 mg/kg (with the exception of mice harbouring SFRC01040 tumours, which received doses of 1 mg/kg with the same scheduling). Vehicle for cisplatin, PLD, paclitaxel, vinorelbine and eribulin treatment was Dulbecco’s Phosphate Buffered Saline (DPBS). Electronic calliper measurements of the primary tumour size were taken twice a week until tumours reached 600-700 mm^3^ or when mice reached ethical endpoint. Data collection was conducted using the Studylog LIMS software (Studylog Systems, San Francisco). Graphing and statistical analysis was conducted using the SurvivalVolume package^101^.

Cisplatin *in vivo* response in PDX was assessed as previously described^59^. One hundred days was chosen as a conservative measure to differentiate between cisplatin sensitivity versus resistance for PDX. We defined response as being “cisplatin sensitive” if the average PDX tumour volume of the recipient mice underwent initial tumour regression with complete remission (CR, defined as tumour volume < 50 mm^3^) or partial remission (PR, defined as reduction in tumour volume of >30% from baseline) followed by progressive disease (PD, an increase in tumour volume of >20% from 200 mm^3^ or nadir post-treatment, if nadir ≥ 200 mm^3^) occurring ≥ 100 days from start of treatment; “cisplatin resistant” if initial tumour regression (CR or PR) or stable disease (SD) was followed by PD within 100 days; or “cisplatin refractory” if three or more mice bearing that PDX had tumours which failed to respond (no CR, PR or SD) during cisplatin treatment (day 1-18).

Time to progression (TTP or PD), time to harvest (TTH), and treatment responses are as defined previously^59^. Stable disease (SD) was achieved if TTP for the treatment group was at least twice has long as TTP for corresponding vehicle treated group.

### Generation of cell lines

An OCS GEMM cell line was generated from a T1 OCS GEMM tumour. Briefly, the tumour was manually minced into a slurry using two scalpel blades and resuspended in DMEM/F-12 GlutaMAX medium (Gibco) supplemented with 10% fetal calf serum (FCS). Cell fragments were subsequently plated on 0.1% gelatin coated plate and passaged aggressively within 3-4 days to retain viable malignant adherent cells until a stable cell line was obtained at p12 onward. Cell identity was confirmed by genotyping (as for GEMM tumours). OCS GEMM cells were grown in DMEM/F-12 GlutaMAX medium (Gibco) supplemented with 10% FCS, 50 ng/mL EGF and 1 μg/mL hydrocortisone in 5% CO_2_ at 37°C.

### Adhesion, invasion assays and 3D growth assays

Adhesion assays were carried out in 96-well plates pre-coated with 2% BSA or 20 μg/ml collagen. GEMM cells were pre-treated for a week with DMSO (vehicle control), 0.2 μM cisplatin or 20 nM eribulin. Pre-treated cells were plated at a cell density of 2 × 10^5^ cells/well in triplicate in pre-coated wells and allowed to adhere for 2 hours. Non-adherent cells were aspirated and adherent cells stained with 100 μl of 0.5% crystal violet (Sigma) dissolved in 20% methanol for 15 min at room temperature. Stained cells were solubilised with 50 μl of 0.1 M citrate buffer in 50% methanol. Adherent cells were quantified by measuring absorbance at 595 nm on a Chameleon Luminescence Plate Reader (Noki Technologies). Transwell migration and invasion assays were carried out as previously described^102^. Briefly, 2.5 × 10^5^ pre-treated GEMM cells (as above) were seeded into Matrigel-coated transwells and medium supplemented with 10% FCS placed in the bottom wells to act as a chemoattractant. Parallel assays were carried out in uncoated control transwell inserts to assess cell migration in the absence of extracellular matrix (ECM). 3D growth assays were carried out as previously described^102^. Briefly, wells of a 48-well plate were pre-coated with 1.5 mg/mL collagen (Thermo Fisher Scientific) in DMEM and incubated at room temperature until collagen became solid. Pre-treated (as above) or untreated GEMM OCS cells were resuspended in 1.5 mg/mL collagen/DMEM, plated at 0.02 × 10^5^ cells/well, and incubated at room temperature until collagen became solid. Medium was added to each well and cells incubated at 37°C/5% CO_2_ for 8-10 days.

### Statistical Analysis

Data was analysed using the Student t-test unless otherwise stated and considered significant when the *p* value was <0.05. All statistical tests were two-sided. Bar graphs represent the mean and standard error across independent experimental repeats unless otherwise stated. Survival analysis was performed using the log rank test on Kaplan-Meier survival function estimates. Statistical significance representations: \**p*<0.05, \*\**p*<0.01, \*\*\**p*<0.001.

### Ethics

Samples for the UK cohort were acquired and utilised under the authority of the NHS Greater Glasgow and Clyde Biorepository (Application Reference 286) following approval by West of Scotland Research Ethics Committee 4 (Reference 10/S0704/60). All animal studies and procedures were approved by the Walter and Eliza Hall Institute of Medical Research (WEHI) Animal Ethics Committee (#2019.024) and performed following guidelines for the welfare and use of animals in cancer research.

## Acknowledgements

We thank S. Stoev, R. Hancock, and K. Barber for technical assistance. We thank Prof Ronny Drapkin, Prof Jane Visvader (and Osaka Bioscience Institute, Japan), and Prof Johannes H. Schulte for kind gifts of the mouse strains used to generate the GEMM. We thank Eisai Co., Ltd. for supply of eribulin. This work was supported by fellowships and grants from the National Health and Medical Research Council (NHMRC Australia; Project grants 1062702 (CLS) and 1104348 (CLS and MJW), Senior Research Fellowship 1116955 (ATP)); the Stafford Fox Medical Research Foundation (CLS, HEB, JB, ATP); the Lorenzo and Pamela Galli Charitable Trust (ATP); Cancer Council Victoria (Sir Edward Dunlop Fellowship in Cancer Research to CLS and Ovarian Cancer Research Grant-in-Aid 1186314 to CIA HEB, CIC CJV and CID GR); the Victorian Cancer Agency (Clinical Fellowships to CLS CRF10-20, CRF16014); CRC for Cancer Therapeutics (PhD top-up scholarship to GH); Research Training Program Scholarship (PhD Scholarship to GH). This work was made possible through the Australian Cancer Research Foundation, the Victorian State Government Operational Infrastructure Support and Australian Government NHMRC IRIISS. The Scottish Genomes Partnership is funded by the Chief Scientist Office of the Scottish Government Health Directorates (grant reference SGP/1) and The Medical Research Council Whole Genome Sequencing for Health and Wealth Initiative. Additional funding was provided by the Medical Research Council (the Glasgow Molecular Pathology Node, grant reference MR/N005813/1), Cancer Research UK (grant references A15973 [IMcN] and A17263 [AVB]), the Wellcome Trust (grant reference 103721/Z/14/Z [AVB]) and the Beatson Cancer Charity (grant reference 15-16-051 [IMcN, PR]). Support was also provided by Ovarian Cancer Action, the Cancer Research UK Centres and Experimental Cancer Medicine Centres at both Glasgow and Imperial and the NIHR Imperial Biomedical Research Centre.

The Australian Ovarian Cancer Study Group was supported by the U.S. Army Medical Research and Materiel Command under DAMD17-01-1-0729, The Cancer Council Victoria, Queensland Cancer Fund, The Cancer Council New South Wales, The Cancer Council South Australia, The Cancer Council Tasmania and The Cancer Foundation of Western Australia (Multi-State Applications 191, 211 and 182) and the National Health and Medical Research Council of Australia (NHMRC; ID199600; ID400413 and ID400281).

The Australian Ovarian Cancer Study gratefully acknowledges additional support from Ovarian Cancer Australia and the Peter MacCallum Foundation. The AOCS also acknowledges the cooperation of the participating institutions in Australia and acknowledges the contribution of the study nurses, research assistants and all clinical and scientific collaborators to the study. The complete AOCS Study Group can be found at www.aocstudy.org. We would like to thank all of the women who participated in these research programs.

## Author contributions

C.L.S., M.J.W., I.A.M., H.E.B, and A.T.P. designed the study, developed methodology, analysed data, wrote the manuscript and supervised the study. G.Y.H. and E.L.K. performed experiments, analysed data, and wrote the manuscript. J.B. analysed data, supervised the study and reviewed the manuscript. E.L., C.J.V. and O.K., developed methodology, performed experiments, analysed data and reviewed the manuscript. D.P.E., R.U.-G., U.-M.B., S.D., G.B. and G.R. performed experiments and reviewed the manuscript. H.B.M. analysed data, wrote and reveiwed the manuscript. P.R., R.M.G. and A.V.B. supervised the study and reviewed the manuscript. S.L.C designed the study, developed methodology, analysed data, supervised the study and reviewed the manuscript. O.McN., A. DeF., J.W. and D.D.B. acquired data or samples, supervised the study and reviewed the manuscript. N.T. acquired data, provided administrative support and reviewed the manuscript. AOCS acquired data and reviewed the manuscript.

